# A choice disinhibition paradigm to reveal neural correlates of behavioral disinhibition in Frontotemporal dementia

**DOI:** 10.64898/2026.09.21.753256

**Authors:** Aya M. Kobeissi, Andrew R. Craig, Wei-Dong Yao

**Affiliations:** Departments of Psychiatry and Behavioral Sciences and Neuroscience and Physiology, State University of New York, Upstate Medical University, Syracuse, 13210, USA; Departments of Psychiatry and Behavioral Sciences, Neuroscience and Physiology, Behavior Analysis Studies, and Pediatrics, State University of New York, Upstate Medical University, Syracuse, 13210, USA

**Keywords:** behavioral disinhibition, self-control, impulsive choice, frontotemporal dementia, prefrontal cortex, mouse behavior

## Abstract

Behavioral disinhibition is a hallmark symptom of behavioral variant frontotemporal dementia (bvFTD). There is no cure or strategy to slow progression, mainly due to little understanding of how disinhibition occurs and the mechanisms underlying self-control. To date, disinhibition has not been demonstrated in an FTD model. We designed a paradigm to assess disinhibition, separating general cognitive capacity, and found that aged mutant mice display increased impulsive choice but have intact cognition, suggestive of early-mid- stage bvFTD-associated symptoms. Further, the neural mechanisms underlying self-control are dissociable from impulsive choice; CaMKIIα- and somatostatin-expressing neurons in the medial prefrontal cortex, but not the classical anterior cingulate cortex, were activated during self-control. Together, we demonstrate behavioral disinhibition in mutant mice and identified specific regions and cell types that underlie self-control. This paradigm is viable for future work dissecting neural circuits underling impulsive choice and self-control behavior in bvFTD and other neurological conditions.

## Introduction

Behavioral disinhibition refers to diminished self-control leading to impulsive and socially inappropriate behavior, often without regard for negative consequences (1). Characterized by loss of behavioral control, disinhibition is a core feature of behavioral variant frontotemporal dementia (bvFTD) that is highly distressing to patients and their close others (1,2). Disinhibition is a complex condition that manifests in several disruptive behaviors, primarily poor risk assessment, impulsive and reckless actions, and socially unacceptable behaviors such as making offensive comments, inappropriate touching, and sudden aggressive episodes (3), which occur in bvFTD (3,4). Current therapeutics have limited efficacy and there is no cure (3,5,6).

FTD is a progressive neurodegenerative disease characterized by atrophy of the frontal and/or temporal lobes (2,3). Several genetic mutations have been identified as causative for FTD (7), leading to dysregulation of different cellular processes. Despite these molecular and genetic heterogeneities, similar symptoms arise suggesting that the same neural circuits may be affected. The mechanisms leading to disinhibitory behaviors in bvFTD are essentially unknown. Further, little is known about the detailed mechanisms underlying self-control and the manifestation of disinhibition, largely due to the complexity of self-control behavior and limited methods to robustly assess these behaviors in bvFTD models.

We sought to develop a strategy to assess behavioral disinhibition in a bvFTD mouse model, using *CaMKIIα;GR_80_* mice (8). A prevalent mutation in FTD is a G_4_C_2_ hexanucleotide repeat expansion in *C9orf72* (9,10), which leads to several pathologies, including the abnormal formation of dipeptide repeat (DPR) proteins (9). *CaMKIIα;GR_80_* mice express poly(GR), a highly toxic DPR protein that disrupts several homeostatic cellular processes (7,8), leading to neurodegeneration (8). *CaMKIIα;GR_80_* mice exhibit reduced social behavior (8), while maintaining intact memory (8), suggesting that *CaMKIIα;GR_80_* mice recapitulate early- mid-stage bvFTD-associated social symptoms.

We developed a paradigm to assess behavioral disinhibition, self-control, and cognitive capacity as well as to elucidate the underlying neural mechanisms. We adapted a delay discounting task (11) that assesses preference for a small reinforcer delivered immediately as opposed to a larger reinforcer delivered after a delay (11,12,13,14). Subjects exhibit impulsive choice when choosing the small and immediate reinforcer more frequently and display self-control when waiting for the large and delayed reinforcer (15,16,17). Therefore, impulsive choice involves instant gratification despite undesirable consequences, entailing a lack of forethought. This strategy, termed the choice disinhibition test, assesses disinhibition that is behaviorally and mechanistically dissociable from general cognition and self-control.

## Methods and Materials

### Mice

Male and female GR_80_ (8) and CaMKIIα-tTA (Jackson Laboratory, #007004) mice were crossed to establish *CaMKIIα;GR_80_* double transgenic (mutant), GR_80_ only (control), CaMKIIα only (control), or a double-negative (no GR_80_ or CaMKIIα; control) mice. Male and female *CaMKIIα;GR_80_* mice and control littermates were used at 9-13 months of age, coinciding with middle-age and when behavioral impairments occur. Male and female 9-12-month-old C57BL/6J mice (Jax #000664) were also used. No sex-dependent differences were detected, therefore data were combined. Equal or near equal numbers of male and female and mutant and control mice were used and were randomly assigned.

Mice were group housed 2-5 mice per cage, under a 14/10 hr light/dark cycle (6:30-20:30), and with *ad libitum* access to standard food chow and water except during behavior training and testing. The vivarium was maintained at 21-23°C and humidity of 30-70%. All mouse studies and experimental procedures were approved by the Institutional Animal Care and Use Committee of the State University of New York Upstate Medical University and conducted in accordance with the National Institutes of Health “Guidelines for the Care and Use of Laboratory Animals”.

### Food restriction

Mice were handled and weighed for 7 days to establish their weight. Food chow consumed was weighed daily to determine the amount food eaten to maintain body weight. For 2 days, mice were given food pellets or sucrose water in their home-cage to promote acclimation and ensure consumption. Mice were gradually food restricted to 80-90% of their original body weight. Food chow was reduced by 0.1-0.2 g per day per mouse till mice reached 80-90% or their original body weight, which occurs ∼7 days after food restriction begins. During training and testing, food rations were given at the end of the training session and were consumed within a few hours (13:00).

### Choice disinhibition test

#### Operant-conditioning chamber and equipment

Mouse modular operant-conditioning chambers (MedAssociates) were used. The ceiling and two walls were composed of plexiglass. Two walls were composed of aluminum, one of which consisted of a central nose-poke (which could be illuminated and contained a photocell for response measurements) and a house light (28-Vdc) located near the top of the chamber. The opposite wall held two retractable lever presses with a cue light (28-Vdc) located above that could be illuminated. Additionally, central food pellets (BioServ) were delivered via a food dispenser (MedAssociates) or liquid delivery port, delivered via a syringe pump (MedAssociates). The floor consisted of stainless-steel bars. Each modular chamber was housed in a dark sound insulating unit with a fan (MedAssociates).

#### Reinforcers

For experiments using food pellets (20 mg Dustless Precision Pellets, Bio-Serv, Flemington, NJ), one food pellet was used for the smaller reinforcer and 4 food pellets were used for the larger reinforcer. For experiments using liquid delivery, a 10% sucrose water solution was used. The larger reinforcer consisted of 80 μL of sucrose water and the smaller reinforcer consisted of 20 μL of sucrose water, except in Fig. 1c where the larger reinforcer consisted of either 100 μL or 60 μL of sucrose water and the smaller reinforcer consisted of either 10 μL or 20 μL of sucrose water. All sucrose solutions were mixed daily.

**Fig. 1.**
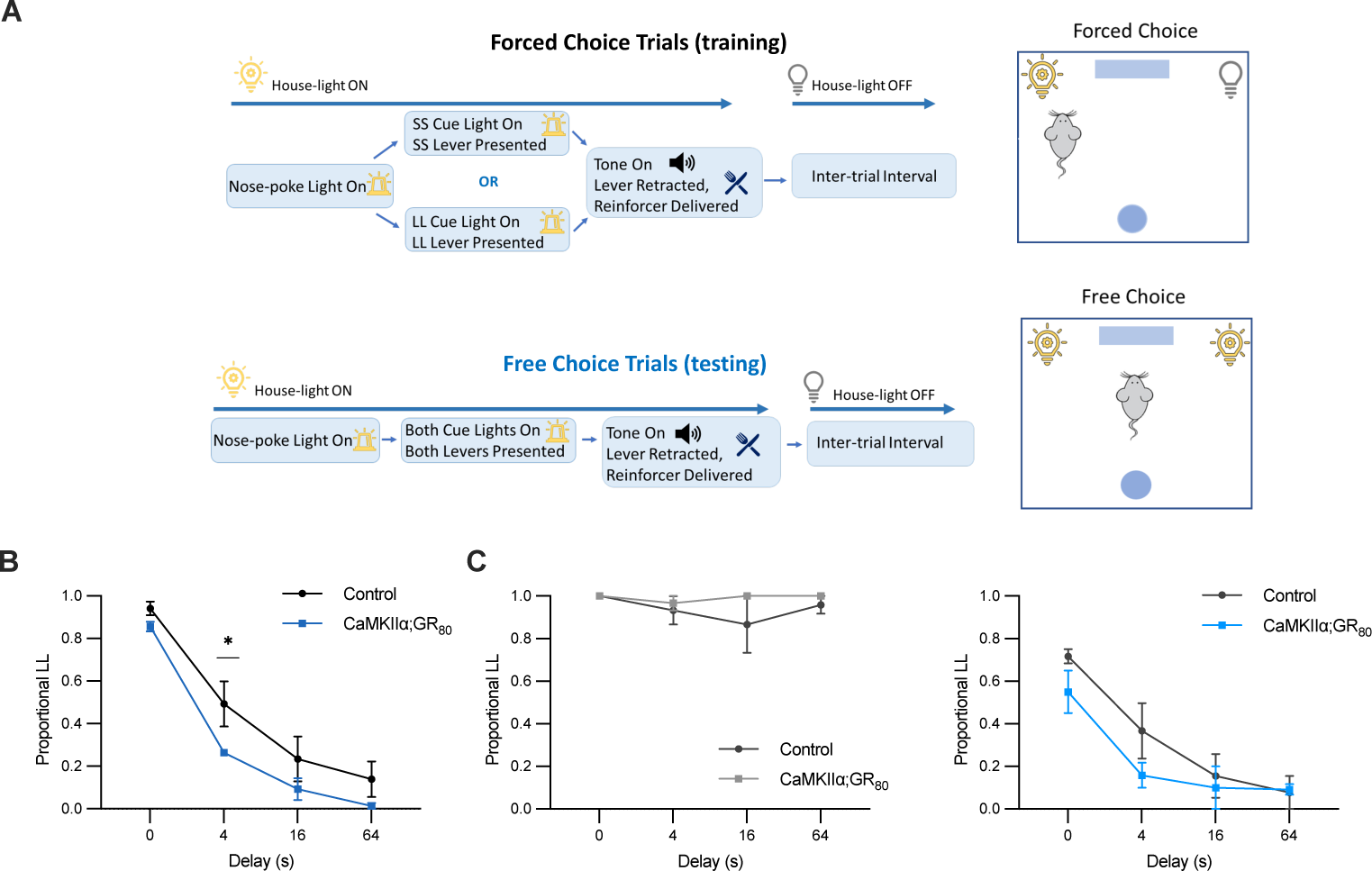
*CaMKIIα;GR_80_* mice exhibit behavioral disinhibition in a choice disinhibition assay. (**A)** Schematic of the forced and free choice trials in the choice inhibition test. LL, larger-later reinforcer. SS, smaller-sooner reinforcer. (**B)** Quantification of the proportion of larger-later reinforcer choice verses small-sooner reinforcer choice, represented as a proportion of the larger-later reinforcer across increasing delays in control and *CaMKIIα;GR_80_* mice during free choice trials. A proportional LL value of 1.0 indicates that the larger-later reinforcer was chosen every time. 80 μL sucrose water was used for larger-later reinforcer and 20 μL sucrose water for the smaller-sooner reinforcer. (**C)** Quantification of proportional LL when sucrose water delivery was largely different (left) between LL (100 μL) and SS (10 μL) or less different (right) between LL (60 μL) and SS (20 μL). In **(A-C)** n= 4 control and n=3 *CaMKIIα;GR_80_* mice. Statistics: **(B-C)** Two-way repeated measures ANOVA with Tukey’s post hoc test. An average of five trials per mouse were used. *p < 0.05, **p < 0.01, ***p < 0.001, ****p < 0.0001. Data are represented as mean ± s.e.m.

#### Pretraining

For experiments involving food pellets, single food pellets were used as reinforcers, and 20 μL of sucrose were used as reinforcers in experiments involving sucrose during pretraining. Mice were first trained to consume reinforcers, which were delivered response independently according to a variable-time 60-s schedule. Magazine training lasted until mice reliably consumed delivered reinforcers (∼five 30-min sessions).

Next, mice were trained to nose poke to produce reinforcers. Here and for the remainder of pretraining, daily sessions lasted 30 min or until 40 reinforcers were earned, whichever occurred first. They were then trained to chain nose poking and lever pressing. Nose pokes produced extension of a lever (left/right counterbalanced) into the chamber, and pressing the lever produced retraction of the lever and delivery of a reinforcer. Nose-poke/lever-press chaining was repeated with whichever lever the mouse was not exposed to initially. The terminal phase of pretraining entailed nose poking pseudorandomly producing the left or right lever with the caveats that the same lever could not be presented more than two times in succession and that both levers were presented an equal number of times during sessions. Mice progressed through operant training when they reliably produced ≥ 30 reinforcers by engaging in the specified operant response (nose poke or chained nose poke and lever press) for three consecutive sessions. The purpose of requiring a nose-poke prior to lever press was to ensure that mice did not become situated in front of a specific lever, thus biasing subsequent assessment of choice. This also assessed their ability to learn the nose-poke plus lever press sequence as well as their motivation to acquire and continue the task (cognitive capacity).

#### Training larger-later and smaller-sooner responses

In a counterbalanced manner among mice, a smaller reinforcer was paired to one lever and was delivered immediately (smaller-sooner) after a lever press (0 s delay). The other lever was paired with a larger reinforcer delivered at progressively increasing delays (larger-later) during each block. Each session consisted of 40 total trials, one session per day, with four 8-trial blocks. Each block consisted of 4 forced trails where only one lever was presented. Mice could not move onto the next phase until a choice was made. Of the forced trials, 2 were designated as smaller-sooner and 2 were designated as larger-later. 4 free choice trials were administered after a forced trial block, where both levers were presented together, and mice could choose which lever to press.

First, no delays were given for the larger-later reinforcer (0 s delay) across all trials within a session. The purpose of this phase was to ensure that mice’s choice making was sensitive to reinforcer magnitude (defined as choosing the larger reinforcer on ≥ 80% of trials). After successful completion, the next set of delays were introduced: block 1- 0 s, block 2- 1 s, block 3- 4 s, and block 4- 8 s; then block 1- 0 s, block 2- 4 s, block 3- 8 s, and block 4- 16 s; and last block 1- 0 s, block 2- 4 s, block 3- 16 s, and block 4- 64s. Sessions occurred daily.

#### Delay discounting and impulsive choice testing

For testing, the same procedure was used as the training LL and SS phase, when only the following delays were used: block 1- 0 s, block 2- 4 s, block 3- 16 s, and block 4- 64s. Sessions occurred daily until mice produced consistent responses, at which the last 5 sessions were used for data analyses.

#### Delay discounting for c-Fos mapping

Delay discounting was conducted as normal, except that block 4 (64 s delay) was removed and sessions occurred 5 days a week. Rations were increased by 0.2 g on days off. These modifications elicited greater self-control in wild-type mice, even at 16 s delays, and were used to assess self-control as mice waited for reinforcers. Sessions occurred until mice produced consistent responses.

#### Behavioral data acquisition and analyses

Five testing sessions (over 5 days) were measured and averaged for data analyses. Proportional larger-later (LL) was calculated as the portion of larger-later choices made out of all free-choice choice opportunities within a trial block, across increasing delays for the larger reinforcer. Behavior data was automatically acquired via responses made in the operant-conditioning chambers linked to MedPC software (MedAssociates, St. Albans, VT).

### Immunohistochemistry, microscopy, and quantification

After the last behavioral testing session, mice rested in their home-cage for 60 min. Mice were deeply anesthetized with isoflurane (20% v/v in proproleyne glycol) in the home-cage. The brain was drop fixed in 4% paraformaldehyde in 1x phosphate-buffered saline (PBS) for 24-48 hrs. Coronal slices were made at 60 μm with a vibratome (Vibratome 1000) and washed in PBS. The slices were permeabilized and blocked for 1.5 hrs at room temperature in 1% Triton X-100 (Sigma Aldrich, St. Louis, MO) and 10% goat serum (Invitrogen, Carlsbad, CA) in PBS. Slices were washed and incubated overnight at 4°C in primary antibody solution composed of 0.1% Triton X-100, 5% goat serum, and rabbit anti-c-Fos (1:1000, Synaptic Systems, Gottingen, Germany) in PBS. One of the following antibodies was also added: mouse anti-CaMKIIα (1:250, Invitrogen), guinea pig anti-somatostatin-28 (1:500, Synaptic Systems), or anti-parvalbumin (1:1000, Invitrogen). Slices were washed in PBS and placed in DAPI (1:1000, Invitrogen) in PBS for 10 min at room temperature, and mounted onto Superfrost Plus slides (Fisher Scientific, Waltham, MA) with Prolong Gold Antifade (Invitrogen).

Images were acquired with a confocal microscope (Leica SP8, Leica Biosystems, Nussloch, Germany) at 20 x magnification and processed using automated mosaic stitch acquisition software (Leica Suite). Images were overlayed with the Mouse Brain Atlas (18) and regional borders of the medial prefrontal cortex (mPFC) and anterior cingulate cortex (ACC) were defined. The following anterior-posterior (relative to Bregma) were used: mPFC, 2.10 mm to 1.54 mm and caudal ACC, 1.42 mm to −0.22 mm. Marker-positive cells were counted manually using LAS X Office software (Leica), within regional bounds. between mice.

For quantifications involving c-Fos, 3 brain slices per mouse were used and the same 3 coordinates were represented for each mouse. The number of marker-positive cells were summed per slice and plotted as a data point. For co-localization analyses, the number of c-Fos-positive cells with a marker of interest was divided by the total number of c-Fos cells.

## Statistical analyses

All data are shown as mean ± s.e.m. c-Fos and some behavior data were statistically analyzed with two-sided unpaired Student’s t-tests for between-group comparison of two variables. Behavior data from delay discounting training and testing were analyzed with two-way repeated measures ANOVA with Tukey’s post-hoc test. The significance threshold was set to α = 0.05; *p < 0.05, **p < 0.01, ***p < 0.001, and ****p < 0.0001. All statistical tests were conducted using Prism 9.0 (Graphpad Software, La Jolla, CA). No mouse was excluded. Experiments were randomized and not performed blind. All behavior data were acquired and analyzed automatically by MedPC5 (MedAssociates) software.

## Results

### *CaMKIIα;GR_80_* mice exhibit behavioral disinhibition in a choice disinhibition assay

We developed a behavioral paradigm that captures and separates impulsive decision making and self-control. We also assessed cognitive capacity, since cognition is typically spared in early-mid-stage bvFTD and could potentially be a confounding variable in the evaluation of impulsive choice. We established the choice disinhibition test, adapted from a delay discounting paradigm (11), where we trained *CaMKIIα;GR_80_* mice and their control littermates to lever press for a reinforcing sucrose water reinforcer (Figure 1a). *CaMKIIα;GR_80_* mice almost exclusively chose the larger reinforcer when delivered immediately, similar to control mice (p=0.9908; F_2,3_=4.130, p=0.0839) (Figure 1b; Figure S1a). As we introduced increasing delays between a lever press and reinforcer delivery, mice chose the larger and delayed (larger-later) reinforcer less often compared to the smaller and immediate (smaller-sooner) reinforcer, indicating the presence of delay discounting (F_3,20_=48.70, p < 0.001) (Figure 1b). *CaMKIIα;GR_80_* mice chose the larger-later reinforcer less frequently compared to controls, suggesting that *CaMKIIα;GR_80_* mice display impulsive choice (F_1,20_=7.556, p=0.0124) (Figure 1b). Impulsive choice in *CaMKIIα;GR_80_* mice was not due to an inability to discriminate between reinforcer magnitude, as *CaMKIIα;GR_80_* and control mice were able to discriminate between reinforcer sizes that were either significantly different from one another, with the larger reinforcer eliciting wait for delivery (F_1,12_=1.000, p=0.3213), as opposed to when reinforcers were close in magnitude (F_1,12_=2.400, p=0.1443) (Figure 1c). *CaMKIIα;GR_80_* mice did not have issue in learning the action sequences needed to receive a reinforcer (F_2,3_=4.1320, p=0.0839 (a); F_2,15_=2.000, p=0.1617 (b); F_4,12_=0.000, p=1.000 (c)) (Figure S1a-c) nor acquiring the delay discounting behavior (Figure 1b,c). These data suggest that *CaMKIIα;GR_80_* mice exhibit increased impulsive choice but intact cognitive capacity, suggesting selectively impaired self-control.

### Select cell types are activated in the mPFC during self-control

The prefrontal cortex (PFC) has been previously shown to regulate self-control and its loss in humans (1), but the underlying cell types and microcircuits within the PFC were unclear. We conducted a c-Fos brain activity mapping assay to determine the fronto-cortical regions activated during self-control (Figure 2a,b). We modified the choice disinhibition task so that wild-type C57BL/6J mice exhibit self-control during two delay periods (F_3,36_=0.5000, p=0.6793 (a left); F_3,36_=1.100, p=0.3425 (a right); F_3,3_=1.025, p=0.0750 (c)) (Figure 2a; Figure S2a-c). The “with delay” group received four and sixteen second delays for the larger-later reinforcer. The “no delay” group received the same protocol as the “with delay” group, except that there were no delays for the larger reinforcer, thereby removing the need for self-control. We found that the caudal anterior cingulate cortex (ACC), comprising of posterior Cg1 and Cg2 regions, did not show differential c-Fos expression between groups (cCg1 t=0.9600, p=0.8765; Cg2 t=0.6780, p=0.9691) (Figure 2c). However, the mPFC, which is comprised of the rostral ACC (rostral Cg1), prelimbic, and infralimbic areas, had significantly more c-Fos expression in the “with delay” group, suggesting that the mPFC exhibits increased cell activation during self-control (rCg1 t=3.711, p=0.0040; PL t=2.711, p=0.0219; IL t=3.678, p=0.0062) (Figure 2c). To determine the specific cell types that were activated during self-control, we assessed co-localization of c-Fos with three major cell types in the mPFC (Figure 2d-f). We found that c-Fos was co-localized with CaMKIIα- (rCg1 F_8,8_=8.711, p=0.0002; PL F_8,8_=1.276, p<0.0001; IL F_7,8_=8.023, p<0.0001) (Figure 2d) and somatostatin-containing (rCg1 F_8,8_=1.507, p=0.0064; PL F_8,7_=1.746, p=0.0004; IL F_7,5_=1.126, p=0.0015) (Figure 2e) cells. However, c-Fos was essentially not co-localized with parvalbumin-containing cells (rCg1 F_7,8_=1.334, p=0.8699; PL F_7,8_=1.005, p=0.8927; IL F_7,8_=6.021, p=0.5856) (Figure 2f). Together, cell types within the mPFC are differentially activated during self-control.

**Fig. 2.**
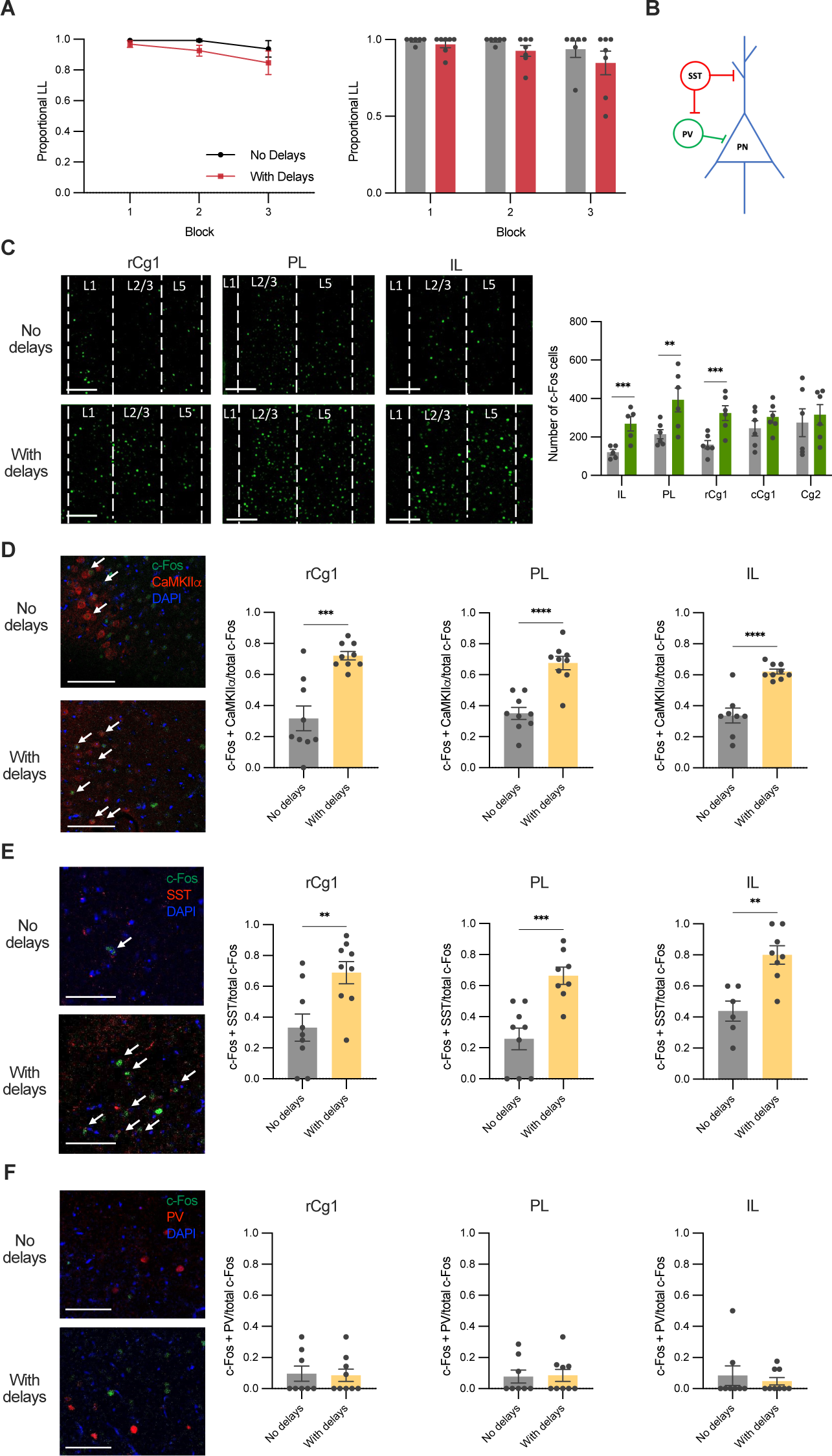
Select cell types are activated in the mPFC during self-control. **(A)** Proportional LL curve (left) and average and individual data points (right) across three blocks during free choice trials. The no delay group did not receive any delays with the small or large reinforcers. The with delay group did not receive delays with the small reinforcers or the large reinforcer in block one. The with delay group received a 4 second delay between lever press and the large reinforcer in block two and a 16 second delay between lever press and the large reinforcer in block three. **(B)** Schematic of a microcircuit in the mPFC. PN, pyramidal neuron. PV, parvalbumin interneuron. SST, somatostatin interneuron. **(C)** Representative photographs (left) of c-Fos cells (green) in the rostral ACC (rCg1), prelimbic area (PL), and infralimbic area (IL) and quantification (right) of c-Fos-containing cells in the rCg1, PL, and IL, caudal Cg1 (cCg1), and Cg2 regions. Scale bar, 150 μm. **(D)** Representative photographs (left) and quantification (right) of cells containing c-Fos (green) and CaMKIIα (red) in no delay and with delay groups. DAPI is shown in blue. Arrows indicate cells with co-localized markers. Scale bar, 75 μm. **(E)** Representative photographs (left) and quantification (right) of cells containing c-Fos (green) and SST (red) in no delay and with delay groups. DAPI is shown in blue. Arrows indicate cells with co-localized markers. Scale bar, 75 μm. **(F)** Representative photographs (left) and quantification (right) of cells containing c-Fos (green) and PV (red) in no delay and with delay groups. DAPI is shown in blue. Arrows indicate cells with co-localized markers. Scale bar, 75 μm. In **(A)** n= 6 mice in the no delay group and n= 7 mice in the with delay group. In **(C-F)** n= 3 representative slices from 3 mice in the no delay group and n= 3 representative slices from 3 mice in the with delay group. A subset of mice from behavior in **(A)** were used for immunohistochemistry in **(C-F)**. Statistics: **(A)** Two-way repeated measures ANOVA with Tukey’s post hoc test. An average of five trials per mouse were used. **(C)** multiple unpaired t-tests with FDR. **(D-F)** Two-tailed unpaired Student’s t-test. *p < 0.05, **p < 0.01, ***p < 0.001, ****p < 0.0001. Data are represented as mean ± s.e.m.

## Discussion

We designed a behavioral strategy utilizing delay discounting to assess self-control behaviors as well as their underlying neural correlates (the choice disinhibition test). We found that *CaMKIIα;GR_80_* mice exhibit increased impulsive choice, which may be modulated by CaMKIIα- and/or somatostatin-expressing neurons. This paradigm robustly captures impulsive choice and cognitive capacity and is effective for delineating the neural circuits and mechanisms underlying loss of self-control. The choice disinhibition test provides a much-needed method for assessing behavioral disinhibition based on decision-making and forethought in animal models.

Impulsivity is multifaceted and encompasses several cognitive and motor variants that are behaviorally and mechanistically distinct (19,20,21). Impulsive choice, which is distinct from impulsive motor actions (22, 23), has been observed in patients with FTD and was correlated with mPFC atrophy (24, 25), suggesting that dysfunction of the PFC may underlie behavioral disinhibition in people with FTD. Studies in animals consistently show that the PFC regulates impulsive choice (26,27,28). Neurons in the prelimbic area of the PFC track impulsive choice and reinforcer subjective value during delay discounting (28). Further, subsets of prelimbic neurons responded to either larger-later reinforcers, or smaller-sooner reinforcers, or both, with the percentage of active neurons shifting across increasing delays in free choice trials (28). In humans, the dorsolateral prefrontal cortex was activated when subjects learned to obtain larger-later reinforcers (29). These results suggest that the PFC regulates impulsive choice, though the PFC microcircuits that drive self-control were unclear.

CaMKIIα-containing pyramidal neurons in the mPFC have long-range afferents to subcortical areas that regulate impulsivity (12,30,31). Inhibition of the mPFC projections to its downstream targets shifts preference to smaller-sooner reinforcers, thereby inducing impulsive choice (27,32), suggesting that the mPFC may exert top-down control over other regions to regulate impulsivity. Since activation of mPFC pyramidal neurons promotes impulse control, it is logical that we observed activation of CaMKIIα neurons but not parvalbumin interneurons (33), which generally exert inhibition onto pyramidal neurons in the mPFC. Somatostatin interneurons play a complex role in terms of activation or inhibition of pyramidal neurons and can directly inhibit pyramidal neurons and/or provide disinhibition of pyramidal neurons via inhibition of other interneurons (34–37). Since we observed activation of somatostatin interneurons, it is possible that pyramidal neurons are activated during self-control and subsequently activate somatostatin interneurons, suggesting that somatostatin activation during self-control might function as feedback inhibition of pyramidal neuron firing. Therefore, somatostatin interneurons may play a role in fine tuning self-control circuits or may promote self-control via activation of CaMKIIα neurons. Future studies are needed to confirm these hypotheses.

We established a choice inhibition test that robustly assesses behavioral disinhibition in a bvFTD mouse model. Future studies utilizing *in vivo* neural activity recordings and circuit manipulations will provide valuable insights into whether and how specific mPFC cells regulate behavioral disinhibition and how these mechanisms are disrupted in bvFTD. The choice disinhibition test provides an important foundation for future mechanistic studies.

## Acknowledgements

We thank Dr. Fen-Biao Gao for providing the *CaMKIIα;GR_80_* mouse model, Ms. Huihui Dai for administrative and research support, and the Yao laboratory for comments. This work was funded by NIH grants MH106489, NS093097, NS122351, and AG082478 (to W.-D.Y) and NIH grant F99NS139550 and the American Federation for Aging Research Kalman/Biology of Aging Scholarship (to A.M.K.).

