## Supplemental figures and legends for "A choice disinhibition paradigm to reveal neural correlates of behavioral disinhibition in Frontotemporal dementia"

**Supplemental Figure Legends**

**Fig. S1|** Characterization of task performance in control and *CaMKIIα;GR₈₀* mice during training or forced choice trials. **(A)** Quantification of proportional LL in forced choices trials during the training phase in control and $CaMKII\alpha;GR_{80}$ mice. **(B)** Number of responses for the central nose-poke and each lever during training in control and $CaMKII\alpha;GR_{80}$ mice. **(C)** Quantification of proportional LL during forced choice trials in control and $CaMKII\alpha;GR_{80}$ mice. In **(A-C)**, n= 4 control and n=3 $CaMKII\alpha;GR_{80}$ mice. Statistics: **(A)** two-tailed unpaired Student's t-test. **(B)** two-way repeated measures ANOVA with Tukey's post hoc test. **(C)** two-way repeated measures ANOVA with Sidak's post hoc test. An average of five trials per mouse were used. *p < 0.05, **p < 0.01, ***p < 0.001, ****p < 0.0001. Data are represented as mean ± s.e.m.

**Fig. S2|** Additional comparisons during the choice inhibition test. **(A)** Quantification of proportional LL comparing control mice from the $CaMKII\alpha;GR_{80}$ line compared to C57BL/6J (left) and comparing male and female mice (right). **(B-C)** Comparison of responses made during training for food pellets or sucrose water in control mice. **(B)** Number of nose-poke and lever responses for food pellets (left) or sucrose water (right). **(C)** Average number of lever presses made during free choice trials for food pellets verses sucrose water in control mice. In **(A, left)** n= 6 C57BL/6J mice and n= 5 control mice. **(A, right),** n= 3 male and n= 8 female mice. b, n= 5 mice. c, n= 4 mice, within subject for food pellet and sucrose water responses. Statistics: **(A)** two-way repeated measures ANOVA with Tukey's post hoc test. An average of five trials per mouse were used. **(B)** two-tailed unpaired Student's t-test. **(C)** two-tailed paired Student's t-test. *p < 0.05, **p < 0.01, ***p < 0.001, ****p < 0.0001. Data are represented as mean ± s.e.m.

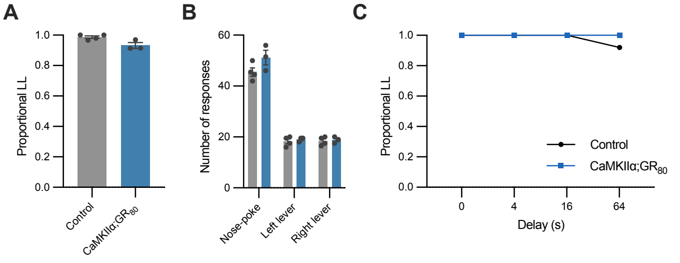

A
1.0
0.8
0.6
0.4
0.2
0.0
Proportional LL
Control
CaMKIIα;GR80
B
60
40
20
0
Number of responses
Nose-poke
Left lever
Right lever
C
1.0
0.8
0.6
0.4
0.2
0.0
Proportional LL
Control
CaMKIIα;GR80
0
4
16
64
Delay (s)

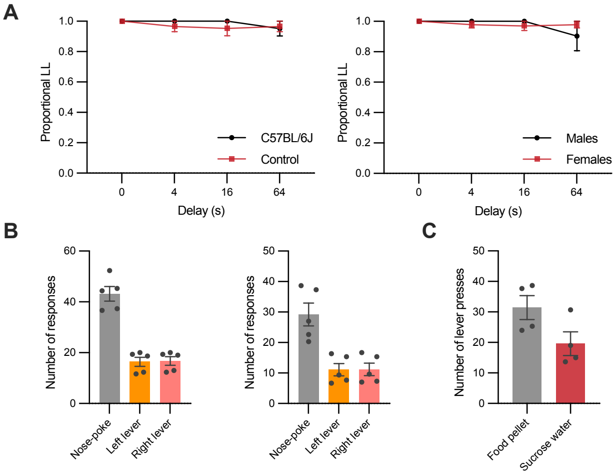
A
Proportional LL
1.0
0.8
0.6
0.4
0.2
0.0
0
4
16
64
Delay (s)
C57BL/6J
Control
Males
Females
B
Number of responses
60
40
20
0
Nose-poke
Left lever
Right lever
50
40
30
20
10
0
C
Number of lever presses
Food pellet
Sucrose water
